# Classification of dog breeds using convolutional neural network models and support vector machine

**DOI:** 10.1101/2023.02.15.528581

**Authors:** Ying Cui, Bixia Tang, Gangao Wu, Lun Li, Xin Zhang, Zhenglin Du, Wenming Zhao

**Affiliations:** National Genomics Data Center, Beijing Institute of Genomics, Chinese Academy of Sciences / China National Center for Bioinformation, Beijing 100101, China; CAS Key Laboratory of Genome Sciences and Information, Beijing Institute of Genomics, Chinese Academy of Sciences, Beijing 100101, China; University of Chinese Academy of Sciences, Beijing 100049, China

**Keywords:** Dog breed classification, Convolutional neural network, Support vector machine, Multi-network integration, Feature selection

## Abstract

Convolutional neural network (CNN) has been widely used for fine-grained image classification, which has proven to be an effective approach for the classification and identification of specific species. For breed classification of dog, there are several proposed methods based on dog images, however, the highest accuracy rate for dogs (about 93%) is still below expectations compared to other animals or plants (more than 95% on birds and more than 97% on flowers). In this study, we used the Stanford Dog Dataset, combined image features from four CNN models, filtered the features using principal component analysis (PCA) and gray wolf optimization algorithm (GWO), and then classified the features with support vector machine (SVM). Eventually, the classification accuracy rate reached 95.24% for 120 breeds and 99.34% for 76 selected breeds, respectively, demonstrating a significant improvement over existing methods using the same Stanford Dog Dataset. It is expected that our proposed method will further serve as a fundamental framework for accurate classification of a wider range of species.

## 1 Introduction

Dog is one of the earliest domestic animals and has a diversity of phenotypes [1, 2]. Currently, there are more than 400 dog breeds worldwide [3], and 283 breeds of them have been registered in AKC (American Kennel Club; https://www.akc.org). Classification of dog breeds is increasingly difficult due to diversities and similarities among dog breeds [4, 5], which extremely dependents on expert experiences. Therefore, identifying the dog breeds easily, accurately and cost-effectively is an interesting challenge for dog breeders, managers or fanciers.

To address this problem, several methods for dog image identification have been proposed, which can be classified into three groups. The first group focuses primarily on training geometric features from dog face images and classifying the features by machine learning method, such as principal component analysis (PCA) [6–8]. However, some of these methods were tested on limited dog breeds (35 breeds) [6, 8] while another one used a large number of dog breeds but the accuracy was not good enough (67% for 133 breeds) [7]. The second group is based on convolutional neural network (CNN) [9–15], which typically employs a single CNN model for Stanford Dog Dataset with most accuracy rate up to 80%. The third group is a combination of CNN and machine learning, in which several studies that focused on improving CNN models [16–19] delivered a better accuracy rate (up to 90%) than single CNN models. In the applications of birds, cats and sheep, these kinds of models archived a classification rate of 95% [20, 21], 80% [22–24] and 85% [25–27], respectively.

Compared to the above methods, the combination of CNN and support vector machine (SVM) used for flower identification has a satisfied accuracy rate of 97% [28, 29]. However, in dog image classification, whether the accuracy of classification can be improved remains to be further explored. Therefore, we initiated this study to integrate multiple CNN networks and machine learning methods, combined with dimension-reduced and feature selection processing, to achieve better identification efficiency.

## 2 Materials and Methods

### 2.1 Dog image dataset

We used the Stanford Dog Dataset [30] to evaluate our model, which is called DataSet1 as follows for convenience. The DataSet1 includes 20,580 annotated dog images from 120 breeds, and each breed has about 180 images. Sample images are shown in Fig. 1.

**Fig. 1.**
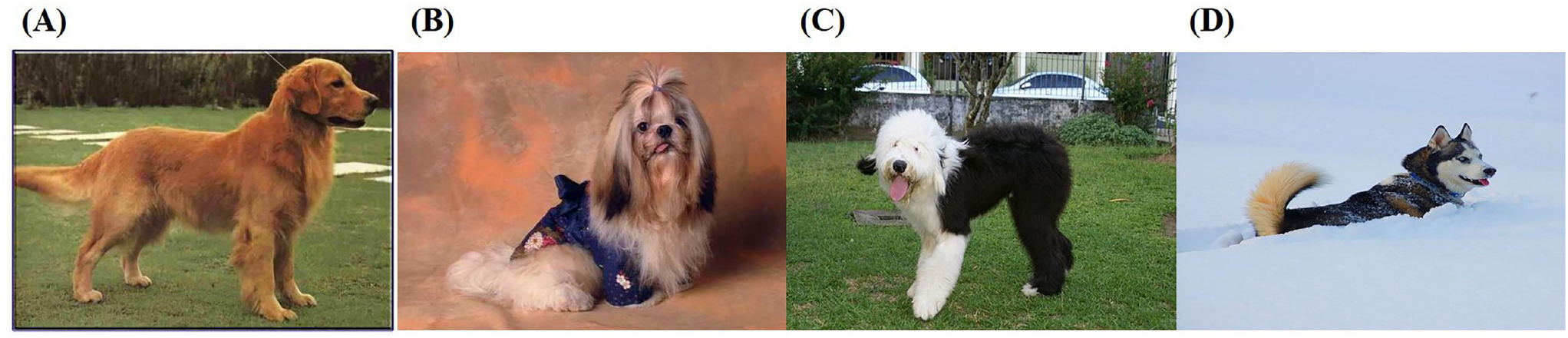
Sample images of Stanford Dog Dataset. (A) golden retriever, (B) Shih-Tzu, (C) Old English sheepdog, (D) Eskimo dog

The distribution of the number of each breed is shown in Fig. 2. Among these 120 breeds, Redbone has the lowest number of images with 148 and Maltese has the largest number of images with 252. In order to eliminate impact of uneven data distribution on accuracy, 120 images were manually filtered for each breed according to image size, aspect ratio and background ratio to form a new dataset named DataSet2. In addition, preprocessing operations were performed on the images, including image size resetting, center cropping, and normalization. The reset image size was set according to requirements of different networks. Both datasets are divided into training and testing set according to the ratio of 8:2, and trained separately.

**Fig. 2.**
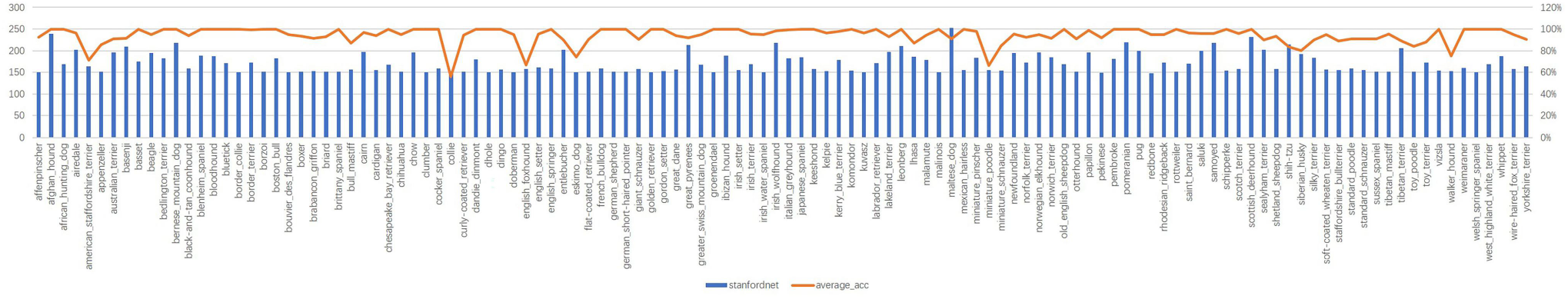
The image distribution and average accuracy of Stanford Dog Dataset by using our best model. The blue colored bars represent the distribution of images of each breed for Stanford Dog Dataset. The orange-colored line represents the average accuracy of each breed using our proposed model.

### 2.2 Proposed model

The model we proposed is shown in Fig. 3, which consists of three main steps.

**Fig. 3.**
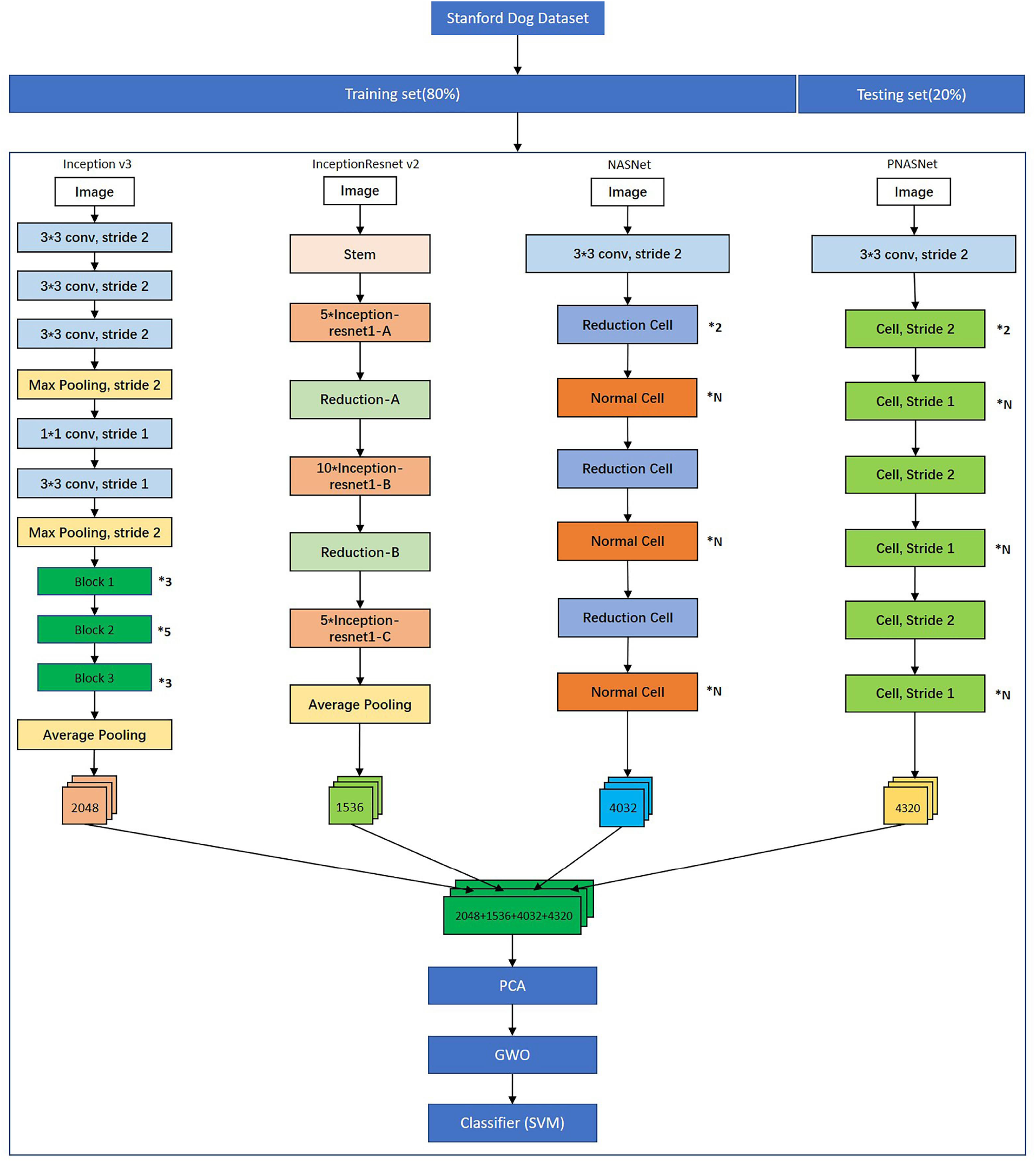
The architecture of proposed model. The Stanford Dog Dataset including Dataset1 and Dataset2 are divided into a training set and a testing set according to 8:2 ratio. Four CNN models including Inception V3, InceptionResNet V2, NASNet and PNASNet are fine-tuned using transfer learning approach and the features before fully connected layer are extracted respectively. 2048 represents the number of extracted features before fully connected layer of Inception V3. 1536 represents the number of extracted features before fully connected layer of InceptionResNet V2. 4032 represents the number of extracted features before fully connected layer of NASNet and 4320 represents the number of extracted features before fully connected layer of PNASNet. And then these features are concatenated and flattened. PCA is used to reduce the feature size and GWO is used to select the specific features. Finally, SVM is used to measure the classification performance.

Firstly, we use Inception V3, InceptionResNet V2, NASNet and PNASNet, which are fine-tuning independent models and have high performance in ImageNet competitions [31–35] to extract features before fully connected layers respectively. Inception V3 is a part of GoogLeNet, which is a deep neural network model based on the Inception module launched by Google. Inception is used to assemble multiple convolution or pooling operations together into a network module [31]. InceptionResNet V2 is an improvement over Inception V3 by introducing residual connections, which reduces computational costs and speeds up network training [33]. NASNet is a new architecture proposed by Google in 2018, which combines features learned in ImageNet classification with the Faster-RCNN framework to exceed the best predictive performance of previously released COCO object detection task. The mean accuracy rate (mAP) of the model was 43.1%, which has 4% improvement over the published best results [34]. PNASNet is an improved model based on NASNet, and proposes a search strategy of Sequential model-based optimization (SMBO). Compared with NASNet, PNASNet is 5 times more efficient, and significantly reduces the requirements of computation resources [35].

All CNN models were trained using transfer learning [36] approach in this study. The size of each input image was set to 299 × 299 pixels for Inception V3 and InceptionResNet V2, and 331 × 331 pixels for NASNet and PNASNet. The four CNN models were trained with 50 epochs respectively using Stochastic Gradient Descent (SGD) with the momentum of 0.9, a mini-batch size of 16 and a fixed learning rate of 0.001. The relevant parameters are shown in Table 1. At the end of transfer learning, the features before fully connection layer of Inception V3, InceptionResNet V2, NASNet and PNASNet were extracted, then we got 2,048, 1,536, 4,032 and 4,320 features respectively.

**Table 1.** Parameter values of CNN architectures

| Software used | Model | Image Size | Optimization | Momentum | Mini Batch | Learning Rate |
| --- | --- | --- | --- | --- | --- | --- |
| Anaconda Python 3.7<br>PyTorch v1 | Inception V3 | 299*299 | SGD | 0.9 | 16 | 0.001 |
|  | InceptionResnet V2 |  |  |  |  |  |
|  | NASNet | 331*331 |  |  |  |  |
|  | PNASNet |  |  |  |  |  |

Next, the features of above four models extracted were combined together, and then filtered using feature selection methods including PCA and gray wolf optimization algorithm (GWO) [37]to obtain specific features. Feature selection can help to improve the accuracy of classification [38]. PCA is a linear dimensionality reduction method, which can reduce the storage space and improve the transmission efficiency. GWO is a meta-heuristic algorithm inspired by gray wolves in nature which mimics their leadership hierarchy and hunting process. GWO designs four agents for simulating the leadership hierarchy, which, from high to low, are gray wolf *α, β, δ* and ω [37]. The description of used parameters for PCA and GWO can be seen from supplemental Table 1.

Finally, SVM was adopted to classify the above selected features, which was originally designed for binary classification problems. When dealing with multi-class problems, it is necessary to construct a suitable multi-class classifier. At present, there are two methods available: 1) Direct method, which can directly modify the objective function and combine the parameter solutions of multiple classification surfaces into an optimization problem. It is simple but difficult to implement because of the high computational complexity and is only suitable for small problems; 2) Indirect method, which mainly realizes multi-classifier by combining multiple binary classifiers. Two common methods are supported, one-against-one and one-against-all [39, 40]. In this study, we used the one-against-one strategy, which was implemented by using Python’s sklearn.svm module, and we selected the RBF kernel function.

We developed the proposed model by integrating above three processes, and realized the combining of CNNs and machine learning methods. The code is implemented in Anaconda Python 3.7, and the deep learning framework is PyTorch (https://pytorch.org/). The whole leaning procedure is runnable on a Linux machine which is equipped with Intel Xeon CPU of 64 GB memory and a Linux machine equipped with NVIDIA GV100GL GPU of 32GB memory.

## 3 Results

### 3.1 The evaluation of single CNN model

The single CNN model can achieve better accuracy but not the best. We used four transfer-learning CNNs to perform the classification, including Inception V3, InceptionResNet V2, NASNet and PNASNet. The results show that NASNet and PNASNet have better classification accuracy than Inception V3 and InceptionResNet V2 (93.03% and 89.64% VS 84.33% and 85.97% in DataSet1, 93.96% and 89.27% VS 86.25% and 84.13% in DataSet2), and NASNet has the highest accuracy in both datasets. The two datasets have similar results and the manually filtered dataset (DataSet2) has little significant advantages. Table 2 summarizes the classification accuracy of each CNN model.

**Table 2.** Result of four models transfer learning

| Models | Test accuracy (Dataset1) | Test accuracy (Dataset2) |
| --- | --- | --- |
| Inception V3 | 84.33% | 86.25% |
| InceptionResNet V2 | 85.97% | 84.13% |
| PNASNet | 89.64% | 89.27% |
| NASNet | 93.03% | **93.96%** |

### 3.2 The evaluation of fusion CNNs and SVM classification

The combined multiple CNNs and SVM improve the classification accuracy compared with the single model above. In our study, the Inception V3, InceptionResnet V2, NASNet and PNASNet fusing with SVM separately achieved a better effect in both datasets. Furthermore, the merged four CNN models fusing with SVM achieved the highest accuracy at 94.1% and 94.9% for 120 breeds in DataSet1 and DataSet2, respectively. Meanwhile, we assembled any two CNNs as mentioned above fusing with SVM, for example, the PNASNet and NASNet (two higher accuracy models) with SVM has higher accuracy (94.2% in DataSet1 and 94.7% in DataSet2) than the combination of Inception V3 and InceptionResNet V2 (two lower accuracy models, 92.6% accuracy in DataSet1 and 93.5% in DataSet2). The results showed that a combination of individual CNN model with high accuracy overall has better performance. Table 3 summarizes the classification accuracy of different models.

**Table 3.** Result of CNN and SVM classification

| Methods | Num of features | Test accuracy (Dataset1) | Test accuracy (Dataset2) |
| --- | --- | --- | --- |
| Inception V3+SVM | 2048 | 89.50% | 91.30% |
| InceptionResNet V2+SVM | 1536 | 92.00% | 92.10% |
| PNASNet + SVM | 4320 | 93.70% | 94.30% |
| NASNet + SVM | 4032 | 93.30% | **94.50%** |
| Inception V3 & InceptionResNet V2+SVM | 3584 | 92.60% | 93.50% |
| NASNet & InceptionResNet V2 + SVM | 5568 | 93.20% | **94.60%** |
| PNASNet & InceptionResNet V2 + SVM | 5856 | 93.80% | 94.20% |
| NASNet & Inception V3 + SVM | 6080 | 93.20% | **94.50%** |
| Inception V3&PNASNet + SVM | 6368 | 93.80% | 94.20% |
| NASNet & PNASNet + SVM | 8352 | **94.20%** | **94.70%** |
| Inception V3 & NASNet & PNASNet & InceptionResNet V2 + SVM | 11936 | **94.10%** | **94.90%** |

### 3.3 The evaluation of feature selection methods

PCA together with feature selection algorithm GWO could improve the accuracy on both datasets, which reached 94.3% on DataSet1 and 95.24% on DataSet2 respectively. Based on the combined four CNN modules above, we used PCA to reduce the features to 5,000, 4,000, 3,000 and 2,000, respectively, then used GWO to select features. In order to evaluate the effective influence of PCA or GWO, we validated the combined patterns of different strategies, for example without PCA or GWO, and the results indicate that GWO has a greater impact on accuracy than PCA (Table 4). The accordingly extracted features of maximum classification accuracy (95.24%) visualized by the t-SNE method is shown in Fig. 4, which clearly shows the different classifications of 120 breeds, and the corresponding confusion matrix (Fig. 5) shows the same pattern.

**Table 4.** Result of four CNN and SVM classification with feature selection

| Methods | Number of features after PCA | Number of features after GWO |  | Test accuracy (Dataset1) | Test accuracy (Dataset2) |
| --- | --- | --- | --- | --- | --- |
|  |  | Dataset1 | Dataset2 |  |  |
| Inception V3 & NASNet & PNASNet & InceptionResNet V2 + SVM | 5000 | - | - | 94.00% | 94.40% |
|  | 4000 | - | - | 94.00% | 94.40% |
|  | 3000 | - | - | 94.00% | 94.40% |
|  | 2000 | - | - | 94.00% | 94.40% |
|  | 5000 | 2508 | 2499 | 94.30% | **95.24%** |
|  | 4000 | 2088 | 2045 | **94.45%** | 95.00% |
|  | 3000 | 1532 | 1483 | 94.23% | 95.07% |
|  | 2000 | 1038 | 1004 | 94.35% | 94.97% |
|  | - | 5991 | 5974 | 94.28% | 95.03% |

**Fig. 4.**
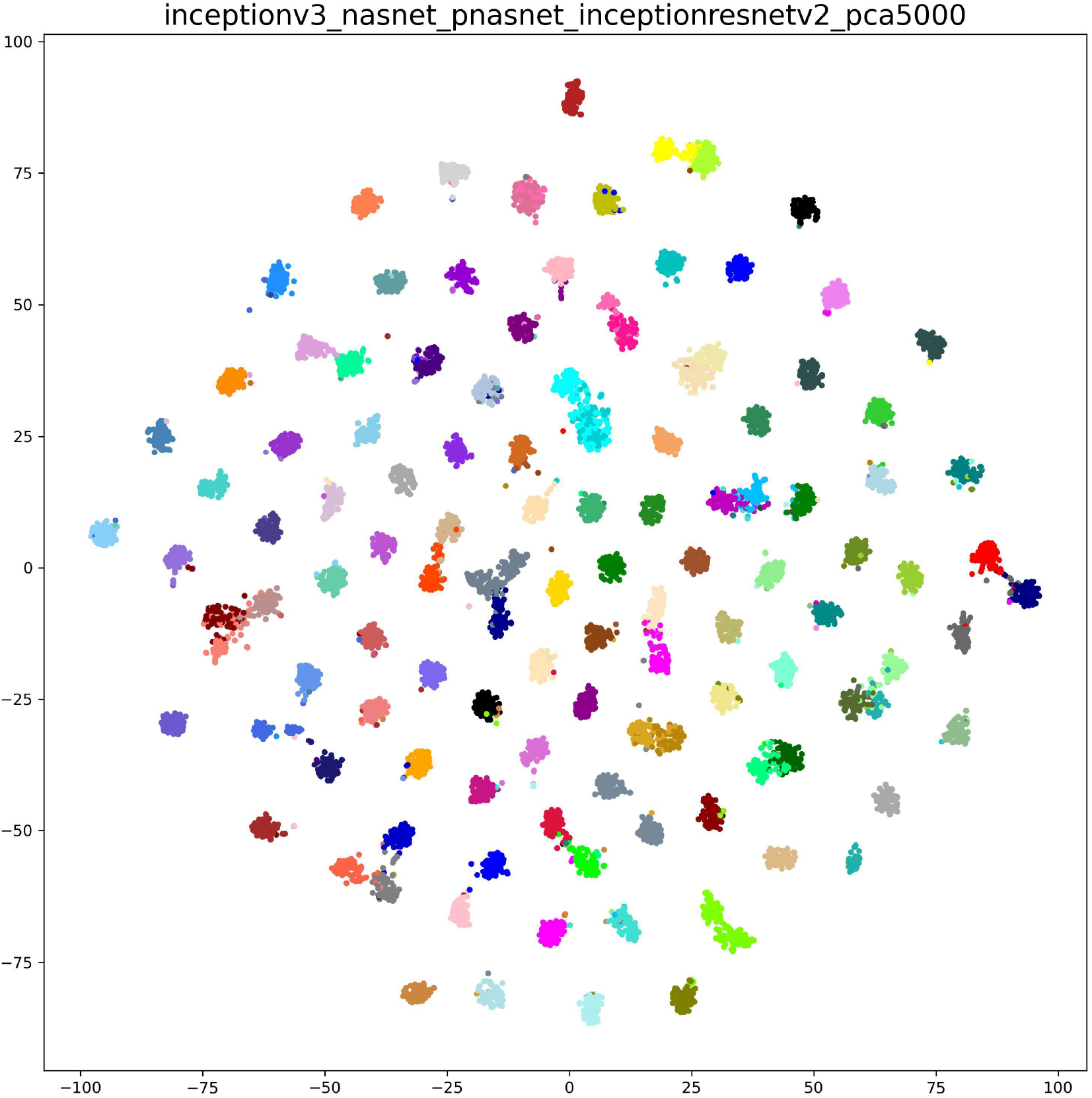
The visualized t-SNE of extracted features of maximum classification accuracy (95.24%) for training dataset of Stanford Dog 120 breeds.

**Fig. 5.**
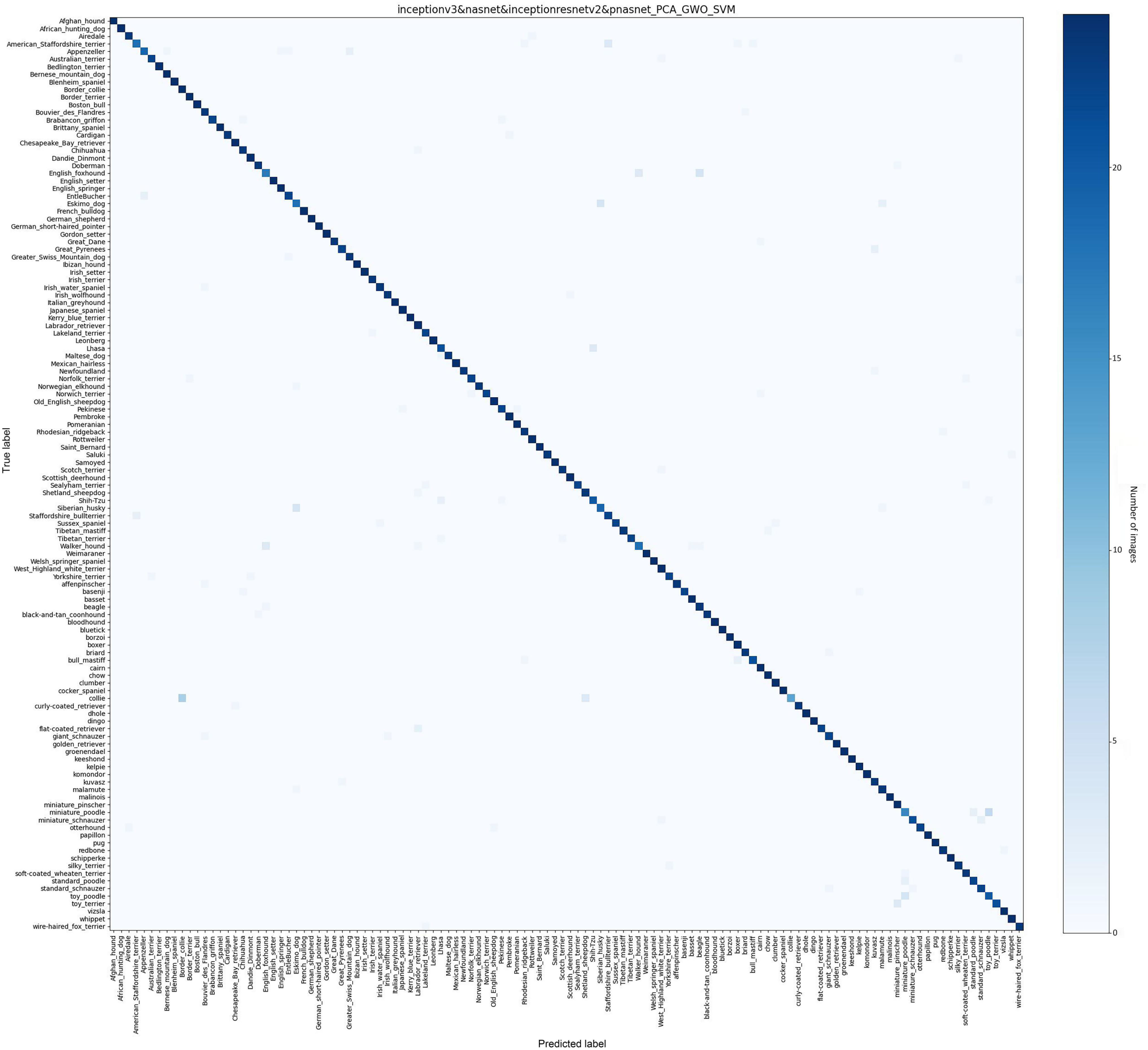
The confusion matrix of maximum classification accuracy (95.24%) for test data of Stanford Dog 120 breeds. The y axis holds the actual label of dog breeds, while the x axis shows the predicted labels.

### 3.4 Summary of results

In total, using the combined method mentioned above for the identification of 120 dog breeds, the average accuracy of each breed for the best model can be seen in Fig. 2 and the accuracy statistics distribution can be seen in Table 5. There are 76 breeds with more than 95% average classification accuracy, of which 48 breeds can be distinguished perfectly with average accuracy of 100%. Three breeds, including English foxhound, Miniature poodle and Collie, have the lowest average accuracy under 70%. We checked out the English foxhound which has average accuracy of 67%. For this breed, it looks more like Beagle and Walker hound which are difficult to be identified even by eyes. In addition, the Stanford images of this breed are different in dog size, photo pose or distance. For the Miniature poodle which has average accuracy 66% and the Collie which has average accuracy 56%, the lower accuracy may be mainly caused by the vast inner difference of the breed such as coat color. We used the top 76 breeds (average accuracy above 95%) to evaluate our model and the accuracy was up to 99.34%.

**Table 5.** Statistics of accuracy distribution of each breed

| Accuracy range | Num of breeds |
| --- | --- |
| 1 | 48 |
| 0.95~0.99 | 28 |
| 0.90~0.95 | 28 |
| 0.80~0.90 | 10 |
| 0.50~0.75 | 6 |

In addition, we developed an online dog image classification tool called DogVC. This tool integrates CNN model, and is available at https://ngdc.cncb.ac.cn/dogvc/. Users can upload a dog image, and then the prediction result will be shown. DogVC was integrated in iDog [41] of NGDC [42], a database of integrating genome, phenotype, disease and variation information of *Canis lupus familiaris*.

## 4 Discussion

Classification of dog breed images is one kind of fine-grained image classification. The proposed model has advantages of comprehensive feature coverage, high-quality feature output and complete automatic implementation. One of the most characteristic features of this study is that we proposed a comprehensive multi-CNN model architecture and conducted experiments of the factors that affect accuracy including model combination, feature size and classified method. We achieved 95.24% accuracy, which is better than other reported deep learning methods using the same Stanford Dog Dataset of 120 breeds regardless of various used hyper parameters (Table 6). Three improvements contribute to the result: 1) Compared to the existing methods that only use single CNN model as backbone model, we combine four CNN models and concatenate the extracted features. Our results show that the fusion of high-performance CNN models have higher accuracy than singe CNN model.

**Table 6.**
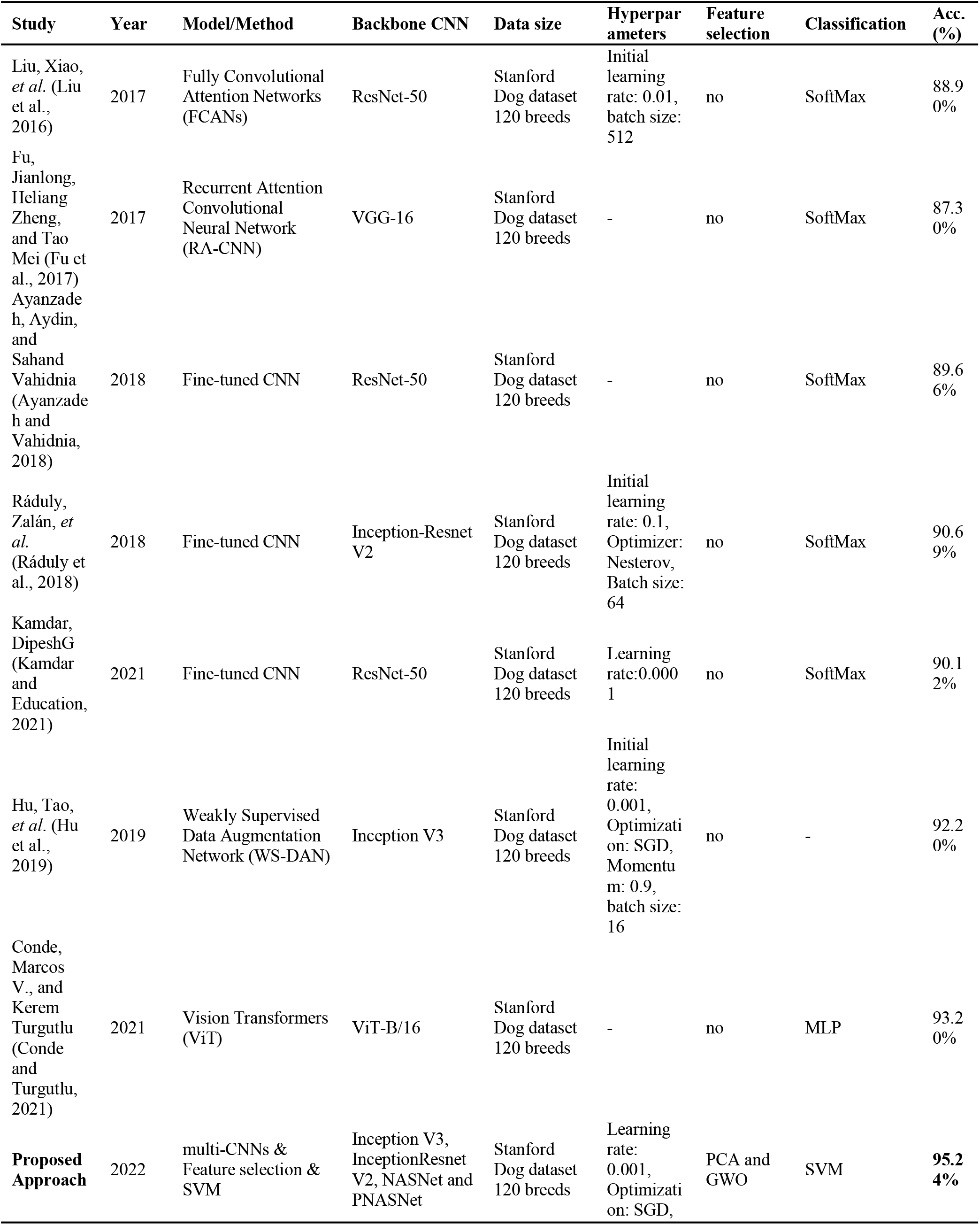
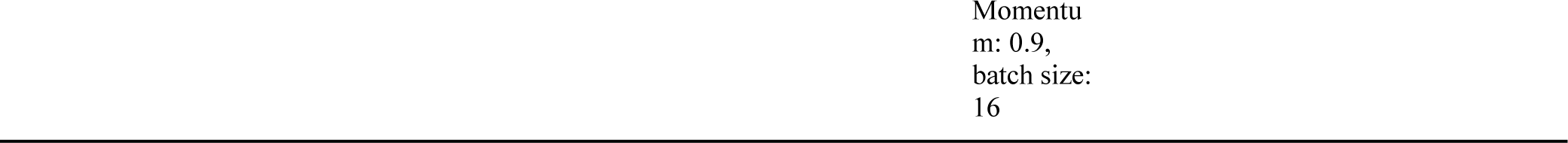
The results of existed studies using Stanford dataset

| Study | Year | Model/Method | Backbone CNN | Data size | Hyperpar<br>ameters | Feature<br>selection | Classification | Acc.<br>(%) |
| --- | --- | --- | --- | --- | --- | --- | --- | --- |
| Liu, Xiao,<br>*et al.* (Liu<br>et al.,<br>2016) | 2017 | Fully Convolutional<br>Attention Networks<br>(FCANs) | ResNet-50 | Stanford<br>Dog dataset<br>120 breeds | Initial<br>learning<br>rate: 0.01,<br>batch size:<br>512 | no | SoftMax | 88.9<br>0% |
| Fu,<br>Jianlong,<br>Heliang<br>Zheng,<br>and Tao<br>Mei (Fu et<br>al., 2017) | 2017 | Recurrent Attention<br>Convolutional<br>Neural Network<br>(RA-CNN) | VGG-16 | Stanford<br>Dog dataset<br>120 breeds | - | no | SoftMax | 87.3<br>0% |
| Ayanzade<br>h, Aydin,<br>and<br>Sahand<br>Vahidnia<br>(Ayanzade<br>h and<br>Vahidnia,<br>2018) | 2018 | Fine-tuned CNN | ResNet-50 | Stanford<br>Dog dataset<br>120 breeds | - | no | SoftMax | 89.6<br>6% |
| Ráduly,<br>Zalán, *et*<br>*al.*<br>(Ráduly et<br>al., 2018) | 2018 | Fine-tuned CNN | Inception-Resnet<br>V2 | Stanford<br>Dog dataset<br>120 breeds | Initial<br>learning<br>rate: 0.1,<br>Optimizer:<br>Nesterov,<br>Batch size:<br>64 | no | SoftMax | 90.6<br>9% |
| Kamdar,<br>DipeshG<br>(Kamdar<br>and<br>Education,<br>2021) | 2021 | Fine-tuned CNN | ResNet-50 | Stanford<br>Dog dataset<br>120 breeds | Learning<br>rate:0.000<br>1 | no | SoftMax | 90.1<br>2% |
| Hu, Tao,<br>*et al.* (Hu<br>et al.,<br>2019) | 2019 | Weakly Supervised<br>Data Augmentation<br>Network (WS-DAN) | Inception V3 | Stanford<br>Dog dataset<br>120 breeds | Initial<br>learning<br>rate:<br>0.001,<br>Optimizati<br>on: SGD,<br>Momentu<br>m: 0.9,<br>batch size:<br>16 | no | - | 92.2<br>0% |
| Conde,<br>Marcos<br>V., and<br>Kerem<br>Turgutlu<br>(Conde<br>and<br>Turgutlu,<br>2021) | 2021 | Vision Transformers<br>(ViT) | ViT-B/16 | Stanford<br>Dog dataset<br>120 breeds | - | no | MLP | 93.2<br>0% |
| **Proposed**<br>**Approach** | 2022 | multi-CNNs &<br>Feature selection &<br>SVM | Inception V3,<br>InceptionResnet<br>V2, NASNet and<br>PNASNet | Stanford<br>Dog dataset<br>120 breeds | Learning<br>rate:<br>0.001,<br>Optimizati<br>on: SGD, | PCA and<br>GWO | SVM | **95.2**<br>**4%** |

2) Existing methods directly use extracted features for classifying, while we use two feature selection methods, PCA and GWO, to obtain improved features. Our results show PCA in combination with GWO has better improvements in accuracy than PCA or GWO alone. 3) Most of the existed methods directly use SoftMax function to classify dog breeds, while we use SVM. Our results show the combination of CNN and SVM can improve the accuracy. The whole results show the feasibility and effectiveness of our proposed model. Meanwhile, the number of images may have no directly effect on breed accuracy (Fig. 2).

Historically, dog image classification has focused on dog facial geometry features such as dog face profile or facial local feature such as ear shape. Due to complexity of localized feature, classified tasks were only trained on limited dog breeds (35 breeds) and/or images (less than 1,000 images) followed by classical machining learning methods such as PCA. Since 2012, the deep learning methods show best-in-class performance in several application tasks, which facilitate the application to dog image classification. By using the Stanford Dog Dataset, single CNN models can classify 120 breeds with the state-of-art accuracy.

Although our method is able to improve the classification accuracy, some limitations also exist. First, since we use the extracted features of four CNN models, the initial feature extracted procedure will take a long time. The merged features from four models will also occupy more resources compared with single CNN model. Second, we only use Stanford Dog Dataset to train our model, and the accuracy as well as generalization of the model used in other datasets such as sheep remain to be verified. In the future, we will focus on further improving the proposed model, and try to use novel CNN networks to extract features such as ViT [19]. In addition, for those lower-accuracy breeds, we will put more efforts into finding underlying causes and potential solutions to improve the overall accuracy. Furthermore, it would be interesting to extend our model to other animal images (such as cats, sheep, birds), so as to explore the scalability and versatility of our model for the task of classification. Finally, a mobile APP should be developed which will widely promote the usage of our model.

## 5 Data and code availability

The Stanford Dog Dataset can be downloaded at http://vision.stanford.edu/aditya86/ImageNetDogs/. DataSet1 (the Stanford Dog Dataset after divided into training set and test set according to 8:2) can b e downloaded at https://download.big.ac.cn/idog/dogvc/dataset_120_raw.zip. DataSet2 (the Stanford Dog Dataset after manually filtered) can be downloaded at https://download.big.ac.cn/idog/dogvc/dataset_120_10.zip. All codes can be found at https://download.big.ac.cn/idog/dogvc/code/ or https://ngdc.cncb.ac.cn/biocode/tools/7319/releases/211 [42]. Besides, all results in this study can be available at https://download.big.ac.cn/idog/dogvc/run_result/.

## 7 Author Contributions

YC programmed the model and performed the analysis. BT designed the model and conducted the analysis. WZ conducted the whole project. YC, BT, XZ and WZ wrote the manuscript. GW participated in Single CNN model testing and topic discussions. LL and ZD also participated in the project discussions and gave valuable advices. All authors contributed to the article and approved the submitted version.

## 9 Acknowledgments

We thank Prof. Guodong Wang and Prof. Yaping Zhang from Kunming Institute of Zoology, Chinese Academy of Sciences, and Prof. Yiming Bao, Prof. Zhang Zhang, Prof. Jingfa Xiao from Beijing Institute of Genomics, Chinese Academy of Sciences / China National Center for Bioinformation for their constructive suggestions.

