## Supplemental Table 1 for "Classification of dog breeds using convolutional neural network models and support vector machine"

Supplymentary Table The parameters of PCA、GWO and SVM

| methods | python function | parameter values |
| --- | --- | --- |
| PCA | sklearn.decomposition.PCA () | n_components(5000,4000,3000,2000) |
| GWO | custom function GWO () | fitness (function to calculate fitness, K-Nearest Neighbor), lb (lower bound, -1), ub (upper bound, 1), dim (seeking range of wolves, here is the dim of input features), Max_iter (number of iterations within algorithm, 5), SearchAgents_no (number of wolves seeking values, 30), NumOfRuns (number of iterations of GWO algorithm, 30), Input feature, input feature label |
| SVM | sklearn.svm.svc () | kernal('rbf'), C (1.0), gamma('auto') |
